## Supplementary material for "*Pseudogymnoascus destructans* transcriptional response to chronic copper stress": SI Figure 1 SI Figure 2

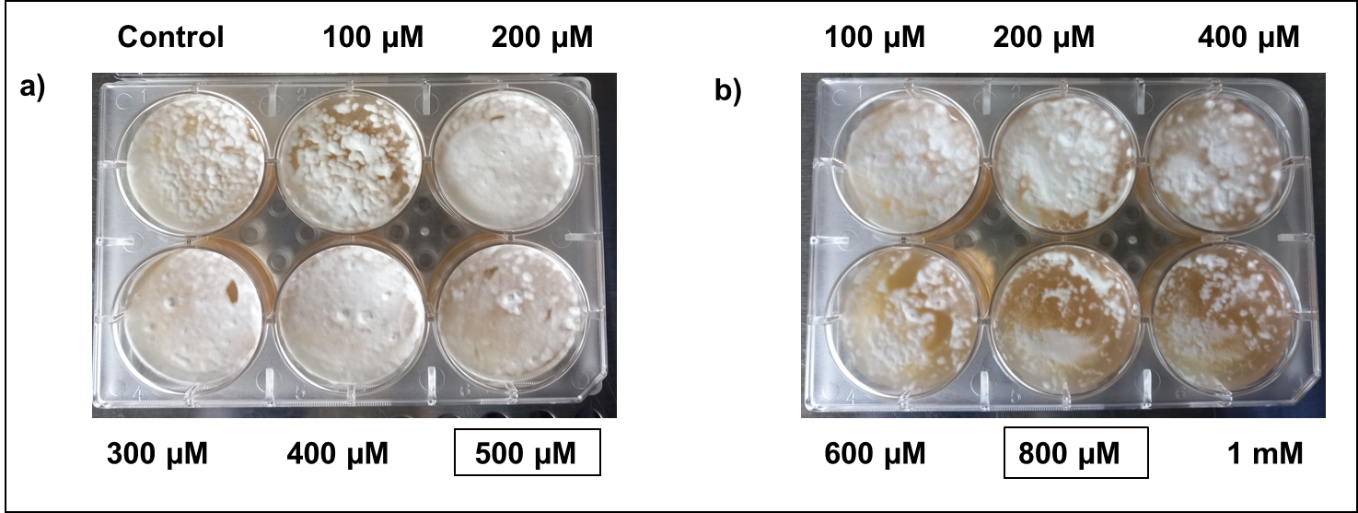

**SI Figure 1.** Growth of *Pseudogymnoasus destuctans* on synthetic (Sc-Ura) plates supplemented with CuSO_4_ or BCS. *P. destuctans* were grown for 6 days under varying concentrations of a) copper and b) BCS stress conditions. Here, we have selected an intermediate concentration of 500 µM CuSO_4_ to stimulate Cu-overload growth conditions and 800 µM copper chelator BCS to stimulate Cu-withholding growth conditions.

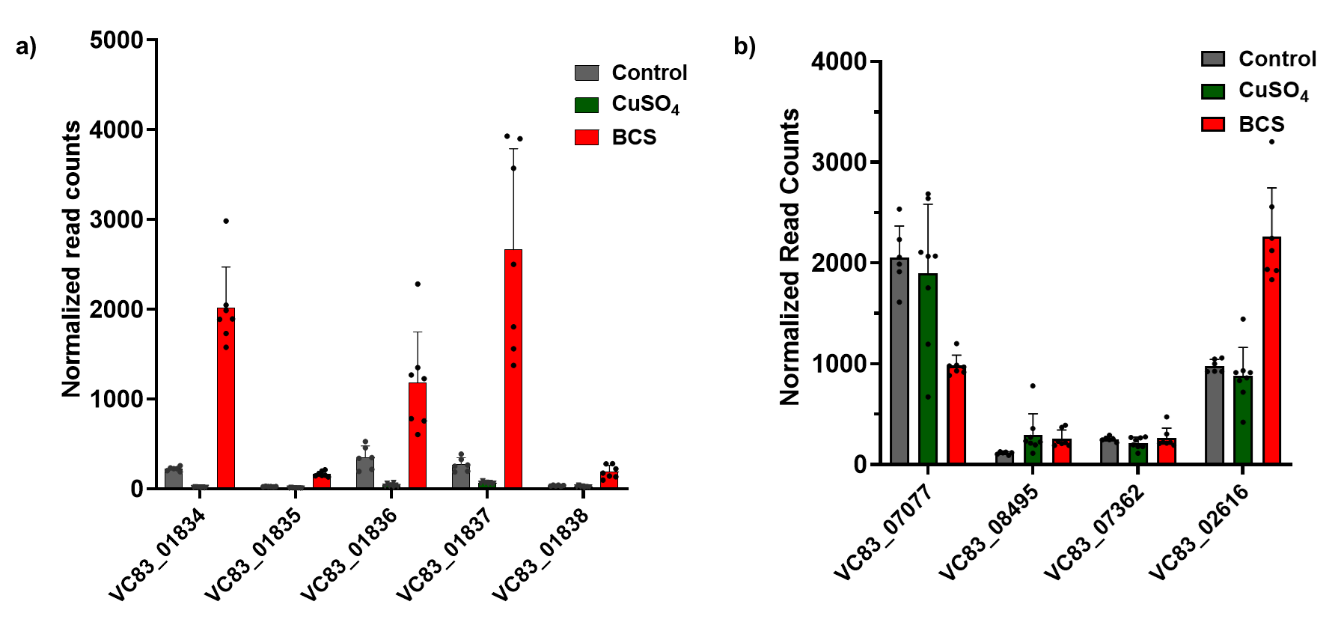

**SI Figure 2.** Normalized read counts for *Pseudogymnoasus destuctans* CRC and SOD genes. a) Copper-responsive gene cluster (CRC). b) Superoxide dismutase (SOD) genes. (Control: n = 6, CuSO₄: n = 8, BCS: n = 7) ****VC83_08495 is multiplied by 10 for graph clarity.**
