## Supplementary material for "*Pseudogymnoascus destructans* transcriptional response to chronic copper stress": SI File 4

***SI File 3.*** *P. destructans* genome encodes for two Mn-SOD enzymes. VC83_07362 contains an N-terminal mitochondrial localization leader sequence while VC83_02616 lacks this peptide motif and may reside in the fungal cytoplasm.

Clustal omega Sequence alignment of VC83_07362 and VC83_02616. Highlighted in yellow is the putative N-terminal mitochondrial targeting sequence for VC83_07362. There is no predicted mitochondrial localization sequence for VC83_02616. Based on these predictions we propose that VC83_02616 may localize in the cytoplasm. The TargetP-2.0 server found at (<https://services.healthtech.dtu.dk/services/TargetP-2.0/>) was used for assignments of Mitochondrial localization signal sequences.

```
VC83_07362      MSATLFRISPAVRSALKAGASKRVARVASTSFVRSKATLPDLQYDYGALEPAISGKIMEL	60
VC83_02616      -------------------------------MSSSKYVLPKLPYAYNALEPYISEQIMTI	29
                                               :  ** .**.* * *.**** ** :** :

VC83_07362      HHSKHHQTYVTSYNAATEQFQAAEAKQDIAAKVALQPLINFHGGGHLNHTLFWENLAPKS	120
VC83_02616      HHSKHHQTYVNNLNIALLSQATAVSTNSLAHQINLQTAIRFNAGGHINHALFWGNLTSAA	89
                **********.. * *  .  :* :.:.:* :: **  *.*:.***:**:*** **:  :

VC83_07362      QGGGRE---PSGALKTAIEDSYGSFIDFQGKFNTALAGIQGSGWAWLVKDNQTGKVLIKT	177
VC83_02616      ETAPSPTSSVAPRLVAALESQWGSVQVFKEKFEAALLAIQGSGWGWLVQDVDTQRLEITT	149
                : .       :  * :*:*..:**.  *: **::** .******.***:* :* :: *.*

VC83_07362      YANQDPVVGQYTPILGVDAWEHAYYLQYENRKAEYFKAIWDVLNWKTAEKRF--------	229
VC83_02616      SKDQDIVPKGKKPLLGIDMWEHAYYLQYLNNKKDYAAGIWNVINWTVVEKRLSTDVDVVF	209
                  :** *    .*:**:* ********* *.* :*  .**:*:**...***:

VC83_07362      ----------	229
VC83_02616      NIVGTLGANL	219
```
