## Supplementary material for "*Pseudogymnoascus destructans* transcriptional response to chronic copper stress": SI Table 1

SI Table 1. Functional Annotation and Expression Profile of 7 Gene Cluster Under Copper Stress

| **Cluster (7 genes)**  **DR C vs BCS and UR C vs Cu** | **Protein Function or Localization** | **Annotation** |
| --- | --- | --- |
| VC83_07925 | ND | Zinc finger |
| VC83_03107 | Secreted | [Hydrophobic surface binding protein A https://doi.org/10.1128/AEM.72.4.2407-2413.2006](https://nam04.safelinks.protection.outlook.com/?url=https%3A%2F%2Fdoi.org%2F10.1128%2FAEM.72.4.2407-2413.2006&data=05%7C02%7Cs_a547%40txstate.edu%7Cf6b6a8293e63485a6b1108dd68a993c0%7Cb19c134a14c94d4caf65c420f94c8cbb%7C0%7C0%7C638781799706790870%7CUnknown%7CTWFpbGZsb3d8eyJFbXB0eU1hcGkiOnRydWUsIlYiOiIwLjAuMDAwMCIsIlAiOiJXaW4zMiIsIkFOIjoiTWFpbCIsIldUIjoyfQ%3D%3D%7C0%7C%7C%7C&sdata=xNcOcYwBACIFv%2FyDrUmg5Y6rCpQrk8URQcdCnKEWpKQ%3D&reserved=0) |
| VC83_07926 | ND | ND |
| VC83_06529 | Secreted | ND |
| VC83_08519 | MFS | Fungal trichothecene efflux pump (TRI12) |
| VC83_05770 | secreted | Aerolisin/ETX pore-forming domain 1 hit (SSF56973) |
| VC83_03991 | Cell membrane | MARVEL domain-containing protein |

* ND not determined
