## Supplementary material for "*Pseudogymnoascus destructans* transcriptional response to chronic copper stress": SI Table 2 SI Table 3

| **Genes** | **Homologs** | | | | **Log_2_FC** | **logCPM** | **Cellular Component** | **Biological process** | **Conserved Protein Domain Family** |
| --- | --- | --- | --- | --- | --- | --- | --- | --- | --- |
|  | ***Saccharomyces cerevisiae* S288C (taxid:559292)** | ***Candida albicans* SC5314 (taxid:237561)** | ***Aspergillus niger* (taxid:5061)** | ***Cryptococcus neoformans* (taxid:5207)** |  |  |  |  |  |
| VC83_07909 | ND | ND | ND | ND | 5.14 | 7.12 | ND | melanin metabolic process | cl09109: Nuclear transport factor 2 |
| VC83_07077 | SOD1 | SOD1 | SOD1 | SOD | -1.07 | 10.53 | ND | superoxide metabolic process | pfam00080: Copper/zinc superoxide dismutase |
| VC83_08495 | ND | SOD5 | GKZ53660.1 | ND | 1.20 | 4.28 | TransMembrane:1 | superoxide metabolic process | pfam00080:Copper/zinc superoxide dismutase |
| VC83_02616 | SOD2 | SOD2, SOD3 | KAI2825965.1 | SOD2 | 1.23 | 10.72 | cytosol | superoxide metabolic process | COG0605:Superoxide dismutase |
| VC83_04976 | RGT2 | HGT12 | GAQ40674.1 | OXC63079.1 | 2.17 | 8.89 | Integral to membrane | carbohydrate transport | pfam00083: Sugar (and other) transporter |
| VC83_01371 | MAL11 | MAL31 | KAI2883056.1 | OXG15676.1 | 2.27 | 7.85 | Membrane | carbohydrate transport | cl26863:Sugar (and other) transporter |
| VC83_05061 | CCP1 | CCP1 | KAI2926762.1 | OXB37400.1 | 1.45 | 8.63 | Mitochondrion | response to oxidative stress | cd00691: ascorbate_peroxidase |
| VC83_00191 | ND | ND | TPR06291.1 | ND | 3.59 | 11.63 | Plasma membrane | copper ion transport | pfam04145:Ctr copper transporter family |
| VC83_01360 | ZRT2 | ZRT1 | GAQ43504.1 | OXG99752.1 | 1.48 | 5.59 | Plasma membrane | metal ion transport | cl00437: ZIP Zinc transporter |
| VC83_00187 | ND | ND | KAL3250585.1 | ND | 1.56 | 3.85 | ND | electron transport | cl12078: Cytochrome P450 |
| VC83_07080 | CCP1 | CCP1 | EHA26204.1 | OXH31575.1 | 1.31 | 7.86 | Cytosol | electron transport | cl00196:Heme-dependent peroxidases |
| VC83_06736 | ND | XP_715252.1 | KAI3006497.1 | OWZ26292.1 | 1.01 | 5.73 | ND | electron transport | cd04730:2-Nitropropane dioxygenase (NPD) |
| VC83_08787 | FRE5 | CFL11 | GKZ79616.1 | OXH01474.1 | 5.37 | 7.88 | Plasma membrane | electron transport | cd06186: NADPH oxidase (NOX) |
| VC83_08524 | DIT2 | ERG5 | KAI3009392.1 | OWZ69196.1 | 3.84 | 6.97 | ND | electron transport | cl12078: Cytochrome P450 |
| VC83_08659 | AIM17 | KHC85949.1 | EHA24645.1 | OWZ80585.1 | 1.54 | 3.34 | Mitochondrion | electron transport | cl26676:Taurine dioxygenase |
| VC83_08976 | BNA4 | BNA4 | GKZ96846.1 | UOH81715.1 | -1.35 | 10.67 | ND | electron transport | cl27552:FAD binding domain |
| VC83_02488 | CBR1 | CBR1 | GAQ41792.1 | OXC64979.1 | 2.16 | 5.24 | ND | electron transport | cl26810:nitrate reductase [NADPH] |
| VC83_02494 | YHB1 | YHB1 | GLA16543.1 | OWT39490.1 | 4.23 | 6.91 | Intracellular anatomical structure | electron transport | cl26811:NAD(P)H-flavin reductase |
| VC83_03586 | BNA4 | BNA4 | KAI2848814.1 | OWZ67419.1 | 1.28 | 7.03 | Mitochondrial outer membrane | electron transport | COG0654:UbiH; 2-polyprenyl-6-methoxyphenol hydroxylase and related FAD-dependent oxidoreductases |
| VC83_03003 | ND | ND | EHA20505.1 | OXC80805.1 | 1.02 | 6.04 | ND | electron transport | cd04730:2-Nitropropane dioxygenase (NPD) |
| VC83_03096 | FRE7 | FRP1 | KAL3253552.1 | OWZ43078.1 | 1.52 | 10.29 | Plasma membrane | electron transport | cd06186:NADPH oxidase (NOX) |
| VC83_03826 | ND | ND | KAI2885967.1 | ND | 1.21 | 6.09 | ND | electron transport | cl00184:Clavaminic acid synthetase (CAS) |
| VC83_04399 | ND | ND | EHA22966.1 | ND | -1.18 | 5.86 | Integral to membrane | electron transport | cd08760: Cyt_b561_FRRS1_like; Eukaryotic cytochrome b(561) |

SI Table 2. Comparative blast analysis of selected significantly regulated genes by Cu-withholding stress (BCS) and their fungal homologs

*ND-Not determined

SI Table 3. Comparative blast analysis of selected significantly regulated genes by Cu-overload stress and their fungal homologs.

| **Genes** | **Homolog** | | | | **Log2FC** | **logCPM** | **Cellular component** | **Biological process** | **Conserved Protein Domain Family** |
| --- | --- | --- | --- | --- | --- | --- | --- | --- | --- |
|  | ***Saccharomyces cerevisiae S288C* (taxid:559292)** | ***Candida albicans SC5314* (taxid:237561)** | ***Aspergillus niger* (taxid:5061)** | ***Cryptococcus neoformans* (taxid:5207)** |  |  |  |  |  |
| VC83_02490 | ND | DAL9 | EHA18141.1 | OXG18647.1 | 3.60 | 4.01 | Integral to membrane | inorganic anion transport | TIGR00886: 2A0108 nitrite extrusion protein (nitrite facilitator) |
| VC83_00102 | ND | ND | GKZ77169.1 | OWZ56854.1 | 1.11 | 4.72 | Cytoplasm | electron transport | cl09933: ACAD Acyl-CoA dehydrogenase |
| VC83_05465 | ERG11 | ERG11 | GKZ67210.1 | OXG23736.1 | 1.64 | 4.39 | Membrane | electron transport | cl12078: p450 Cytochrome P450 |
| VC83_04555 | ND | KHC76775.1 | GLA40930.1 | ND | -1.02 | 1.69 | Membrane | electron transport | ND |
| VC83_08787 | FRE5 | CFL11 | GKZ79616.1 | OWT38774.1 | -1.33 | 2.74 | Plasma membrane | electron transport | cd06186: NADPH oxidase (NOX) |
| VC83_08518 | JLP1 | KHC83894.1 | KAI3021159.1 | UOH83015.1 | 1.84 | 5.65 | Cytoplasm | electron transport | pfam02668: TauD Taurine catabolism dioxygenase |
| VC83_08524 | DIT2 | ERG5 | KAI3009392.1 | OWZ69196.1 | 1.61 | 5.03 | ND | electron transport | cl12078: Cytochrome P450 |
| VC83_08570 | ND | KHC86328.1 | GKZ67210.1 | UOH82104.1 | 1.01 | 2.21 | Membrane | electron transport | cl12078: p450 Cytochrome P450 |
| VC83_08659 | AIM17 | KHC85949.1 | EHA24645.1 | OWZ80585.1 | 1.63 | 3.42 | Mitochondrion | electron transport | cl26676:Taurine dioxygenase |
| VC83_08941 | ERG11 | ALK2 | GKZ77838.1 | OXB38448.1 | 1.00 | 8.13 | ND | electron transport | cl12078: p450 Cytochrome P450 |
| VC83_02428 | FRE7 | FRP1 | EHA27596.1 | OXG26069.1 | 1.26 | 7.00 | Plasma membrane | electron transport | cd06186: NOX_Duox_like_FAD_NADP NADPH oxidase |
| VC83_02488 | CBR1 | CBR1 | GAQ41792.1 | OXC64979.1 | 1.98 | 5.11 | ND | electron transport | cl26810:nitrate reductase [NADPH] |
| VC83_02494 | YHB1 | YHB1 | GLA16543.1 | OWT39490.1 | 1.95 | 4.89 | Intracellular anatomical structure | electron transport | cl26811:NAD(P)H-flavin reductase |
| VC83_03470 | ND | ND | GKZ82666.1 | OWT41612.1 | -1.13 | 6.52 | Membrane | electron transport | cl27552: FAD_binding_3 Superfamily FAD binding domain |
| VC83_03003 | ND | ND | EHA20505.1 | OXC80805.1 | 1.25 | 6.21 | NA | electron transport | cd04730:2-Nitropropane dioxygenase (NPD) |
| VC83_03096 | FRE7 | FRP1 | KAL3253552.1 | OWZ43078.1 | -1.70 | 8.52 | Plasma membrane | electron transport | cd06186:NADPH oxidase (NOX) |
| VC83_04399 | ND | ND | EHA22966.1 | ND | -1.22 | 5.77 | Integral to membrane | electron transport | cd08760: Cyt_b561_FRRS1_like; Eukaryotic cytochrome b(561) |
| VC83_00191 | ND | ND | TPR06291.1 | ND | -4.70 | 7.65 | Plasma membrane | copper ion transport | pfam04145:Ctr copper transporter family |
| VC83_08495 | ND | SOD5 | GKZ53660.1 | ND | 1.52 | 4.52 | TransMembrane:1 | superoxide metabolic process | pfam00080:Copper/zinc superoxide dismutase |
| VC83_00837 | CTS2 | Q5AKZ3.1 | GLA07435.1 | OWZ77110.1 | 1.12 | 4.40 | NA | carbohydrate metabolic process | cl16916: ChtBD1 Hevein or type 1 chitin binding domain |
| VC83_08561 | SGA1 | SGA1 | AIY23067.1 | OXG32870.1 | 1.96 | 7.55 | Vacuole | carbohydrate metabolic process | cd05811: CBM20_glucoamylase Glucoamylase |

*ND-Not determined
